# Bt cotton sustainability needs integration of complementary technologies for the cotton farmers in India

**DOI:** 10.1101/2020.04.30.068262

**Authors:** GT Gujar, CD Mayee, B Choudhary, A Suresh

## Abstract

Biased conclusions of long-term impact of Bt cotton published in Nature Plants March 2020 threatens to derail technological development. We therefore advocate integrating available technologies for sustainability of Bt adoption in India and prospecting for all including biotechnological developments.

---

Bt cotton cultivation continues to draw attention of scientists and others for its impact in India. Latest in this series is the study of Kranthi and Stone^1^ on long-term impact of Bt cotton both prior to and after Bt cotton introduction to elucidate the factors responsible for yield trends. They dismiss the notion that increase in yields of Bt cotton is due to Bt technology and assert importance of other agri-inputs. In their opinion, benefit of Bt technology has been modest and largely ephemeral. Further, Bt cotton adoption is not an indicator of yield trends, but of initial reduction of insecticide use. While admitting the purpose of Bt technology for damage aversion (due to insect attack), authors have focussed more on invalidating the null hypothesis that Bt technology is yield enhancing.

## Role of illegal Bt cotton in %Bt adoption versus Yields

Authors calling the state yield trends with %Bt adoption incongruous set aside a case of Gujarat by justifying presence of illegal Bt cotton contribution for better fit, but dismiss it elsewhere. For instance, the northern states of Punjab, Haryana and Rajasthan showed yield jumps incongruous with Bt cotton adoption. By the time, Bt cotton was approved in these states in 2005, an appreciable area of cotton had in fact come under Bt cotton hybrids to give requisite boost in yields. Peshin *et al*.^2^ reported as much as 22% area under illegal Bt cotton in Punjab by 2004-05, with many Bt cotton hybrids for farmers to grow, and 30.6% higher yield over other varieties. Unfortunately, there is little information on illegal Bt cotton adoption in other two states except for some newspaper reports. As for other locations are concerned, here is an indicator. In 2005, a single district of Yavatmal (with an area of 0.5 million ha cotton) used 90,000 legal Bt packets as against 250, 000 illegal packets^3^. Thus, importance of illegal seeds in the initial years of Bt adoption for yield contribution appears to be under-rated.

## Fallacy that mere share of Bt cotton adoption cannot explain the yield trends

An important point that authors^1^ ignored is the fact that mere quantitative estimate of Bt cotton area adoption is not enough criterion to explain yield trends. It is well known that in the early years of Bt cotton commercialization, approved Bt cotton hybrids were not most suited for various locations. Over the years, as more Bt hybrids were made available, only the best suited were adopted by the farmers in each state^4^. Nevertheless, our correlation analysis of Bt adoption with yields shows a strong coefficient value of 0.61 for data from 2002 to 2018, despite the fact that both Bt adoption and yield data flatten from 2010 and their trends are biphasic in nature. Alternately, we carried out correlation analysis of yield data and %Bt adoption for the year 2007-08 of different states varying from 6.6% to 96.4% which showed coefficient of 0.53, a relatively good relationship between these parameters (Fig. 1). Positive impact of Bt technology on yield increases in Rajasthan and not for Punjab and Haryana using statistically more robust interrupted time series analysis is recently reported (http://ras.org.in/25b7c3bb10f2f5292856121111dad908). This too negates Kranthi and Stone^1^’s observations of increasing yield trends of north Indian states in the early 2000s along with Gujarat dominating the national trends.

**Fig. 1.**
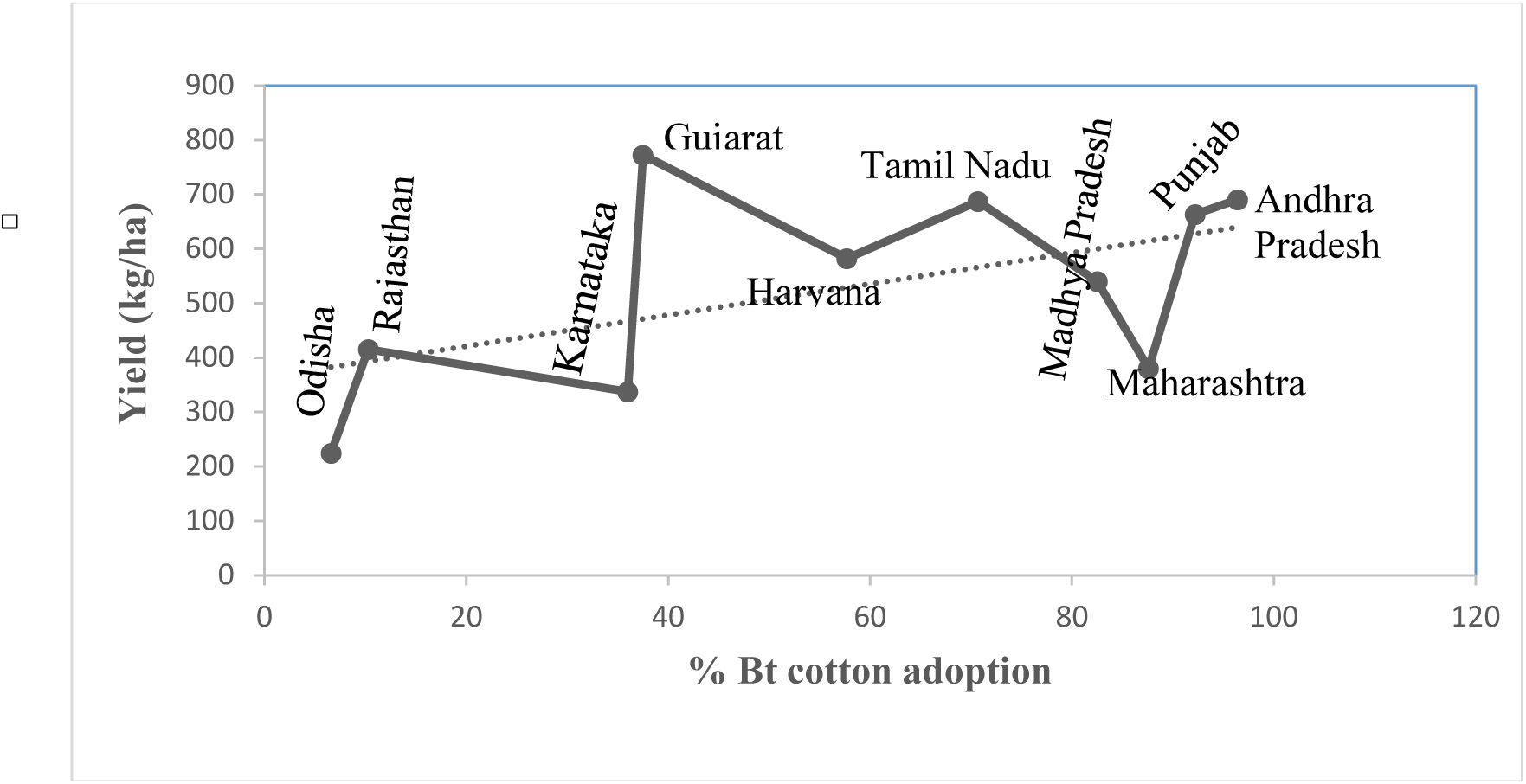
Correlation between % Bt cotton adoption and Yield (kg/ha) of different states at 2007-08.

## Not entirely consequential and dominant effect of fertilizers on % Bt cotton adoption and yields

Another major part of this study is relationship of %Bt adoption and yield trends with fertilizer use over time scale. Fertiliser use efficiency depends upon moisture availability in the soil. And this becomes clear with the data on Rajasthan and Madhya Pradesh wherein fertilizer use over time scale is almost flat, but irrigation seems to explain yield trends.

Maharashtra, a major rainfed cotton producing state in India wherein despite decline in fertilizer use during crucial period of 2002-10, yield increase is more significant than those in later years. Simplistic approach to explain yield association with fertilizer use is pre-mature to ignore these states. The averted boll damage through Bt technology required more nutrients for sustenance of plants. And hence, higher application of fertilizer is to be viewed as a nature of the technology rather than singling out it as a factor responsible for yield increase. Further, of factors that impacted yields adversely during the latter half of Bt cotton phase is declining share of area under irrigation in cotton from 2010-11 onwards (Fig. 2), yet sustaining yields after 2010 should be viewed as a part of interplay of various technological inputs.

**Fig. 2.**
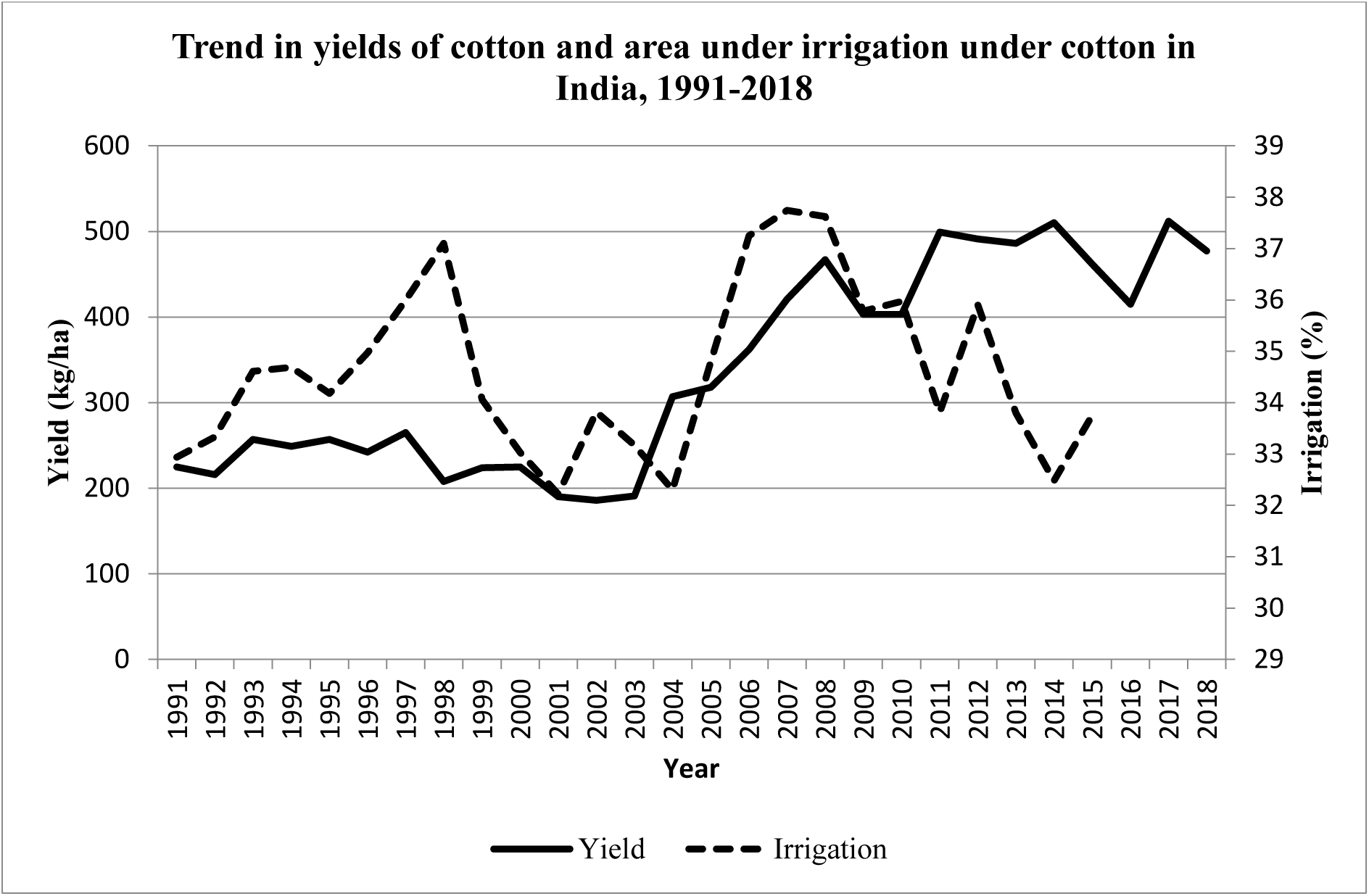
Trends in %irrigation and yield (kg/ha) of cotton.

Another issue worth highlighting is that the imbalanced use of fertilizers that includes nitrogen, phosphorous and potash for most soils, and also zinc, boron and magnesium in some. Farmers use far more nitrogenous fertilizers than the rest of macro-nutrients. Cotton being rainfed crop, fertilizer use efficiency will be limited in view of unpredictability of rains. Using the data provided by Suresh *et al.*^5^ for growth rates (in terms of % per annum) for fertilizer and yield during 2002-03 and 2009-10 (Bt cotton period) for all nine cotton growing states, correlation coefficient was only 0.27 clearly suggesting poor association (Fig. 3). We therefore doubt consequential and dominant effect of fertilizer use on %Bt adoption and yields. On the contrary, imbalanced use of fertilizers has increased incidence of sucking pests, increased costs and affected yields, as reflected in whitefly outbreaks and large scale failure of crop in 2015 in Punjab and other states almost on regular basis.

**Fig. 3.**
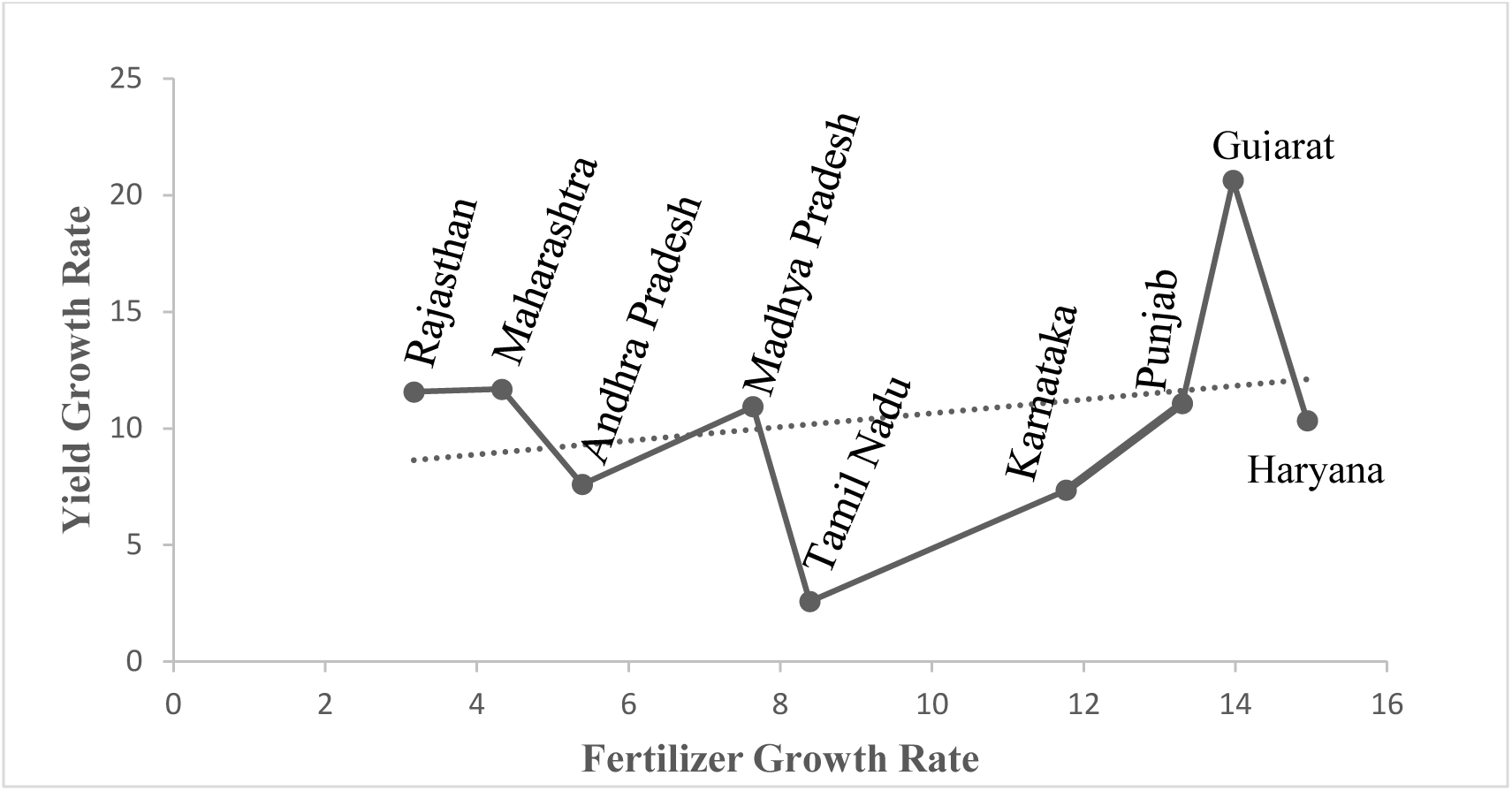
Relationship between growth rate (%/annum) of fertilizer and yield for different states during Bt cotton period of 2002-02 to 2009-10.

## Using wrong unit of insecticide to exaggerate their importance for Bt cotton

Although novel insecticides like spinosad (registered in September 2001) and indoxacarb (registered in July 2001) preceded hardly a year before Bt cotton, their contribution towards jump in yield in 2003 is specially mentioned by authors. We therefore doubt if these novel insecticides were used so extensively as soon as registered on cotton in the country to be solely responsible for yield jump at the national level. Further, the monetary unit of expression for insecticides can never be true indicator of its use. We are not sure if the increasing insecticide cost in US$ reported herein reflects real term costs pegged at base year 1999-2000 accounting for inflation, foreign exchange fluctuations, and overpricing that normally occurs for novel insecticides. On the contrary, Ranganathan and his colleagues (2018)^6^ studying the insecticide use between 2002-03 and 2012-13 reported significant reduction in pesticide costs in real terms (at 2002 prices). Our own analysis of insecticide use in cotton given below in Fig. 4 shows that insecticide use is less than before or at the time of Bt cotton commercialization.

**Fig. 4.**
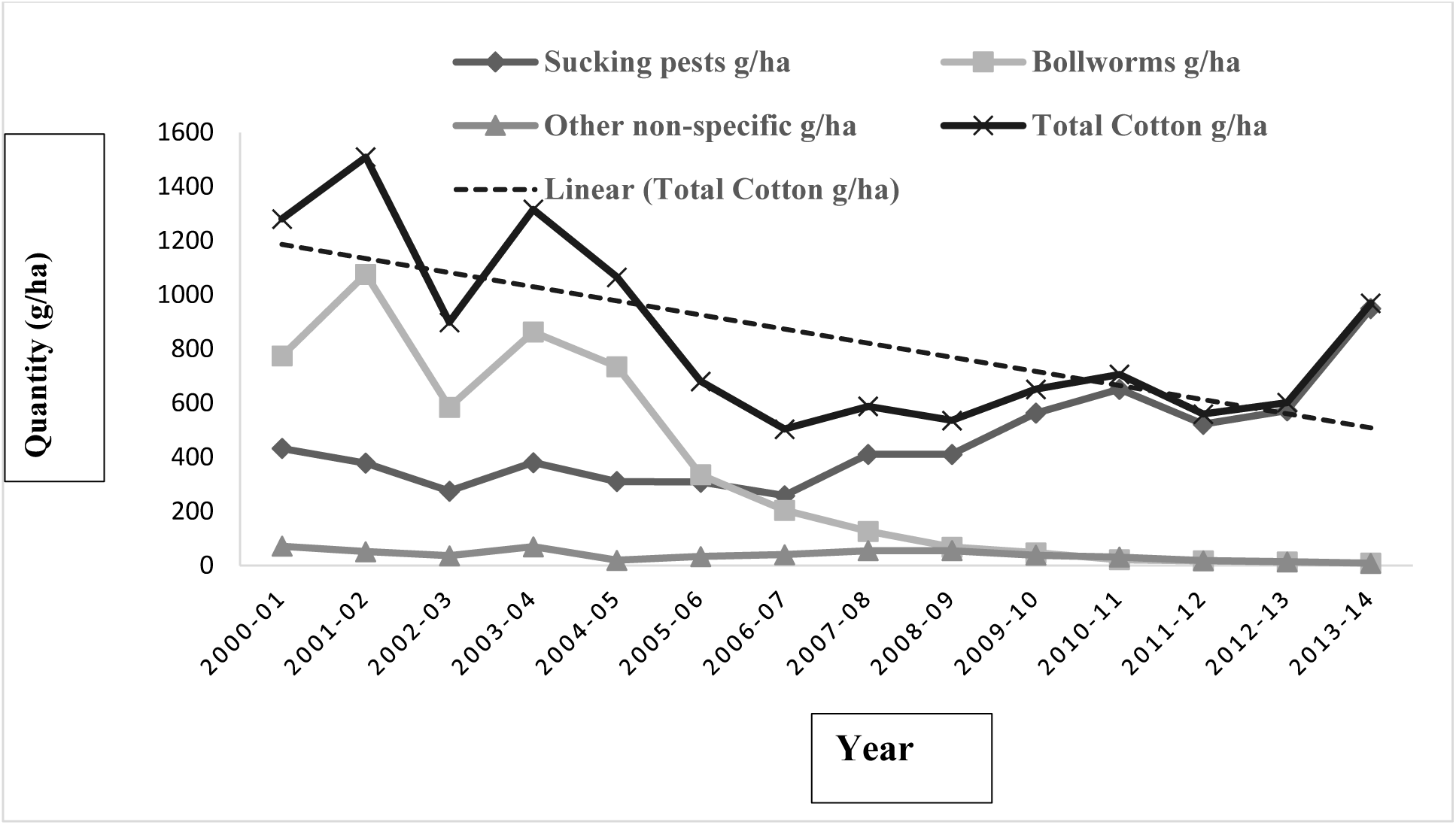
Trends in insecticide use (g/ha) in cotton over 2000-2013. Broken line shows trendline (Based upon the data available from several sources like http://www.cicr.org.in/pdf/pop_art/Fertilizers_and_Bt.pdf, 4, http://ppqs.gov.in/divisions/cib-rc/about-cibrc).

## Ignoring large welfare benefits of Bt technology

This study conspicuously ignores positive shifts that occurred with Bt adoption at reduced real cost of production in all states resulting in large welfare benefits netting out increased cost of cultivation^7^. Fallacy associated with increasing yield trends even before introduction of Bt cotton as claimed by Kranthi and Stone^1^ does not stand scrutiny of increasing yield trends from 2002-03 to 2009-10^7,8^ with some years showing significant yield dips due to drought to bounce back to high of 512 kg/ha in 2017-18. The ignorance of drought impact tends to attribute the yield reduction entirely as the failure of Bt technology.

Failure to quantitatively proportionate yield increases to various farm inputs will often lead to contentious arguments of their importance. As of now correcting imbalances in the current agronomic practices and implementing of integrated pest and resistance management^9,10^ will help to sustaining and even increasing yields. As more than 72% of cotton area belongs to biotech cotton producing about 75% of total production in the world, biotechnology has a definite role to play in the future too and this is reflected in the predominance of Bt cotton area (90-95% of total cotton area) that sustains farmers’ income and hope in India.

